# DTI-ALPS PRIMARILY REFLECTS WHITE MATTER DIFFUSION DISPERSION AND MICROSTRUCTURAL HETEROGENEITY IN NEURODEGENERATION: INSIGHTS FROM MULTI-MODAL MRI

**DOI:** 10.1101/2025.11.12.688102

**Authors:** Luca Sacchi, Giovanni Farruggio, Giorgio Bocca, Tiziana Carandini, Milena De Riz, Laura Ghezzi, Manuela Pintus, Anna Pietroboni, Giacomo Boffa, Elisa Scola, Fabio Triulzi, Marco Bozzali, Daniela Galimberti, Mara Cercignani, Andrea Arighi

**Affiliations:** Department of Biomedical, Surgical and Dental Sciences, University of Milan, Milan, Italy; Neurodegenerative Diseases Unit, Fondazione IRCCS Ca’ Granda, Ospedale Maggiore Policlinico, Milan, Italy; Department of Biomedical Sciences for Health, University of Milan, Milan, Italy; Department of Neuroscience, Rehabilitation, Ophthalmology, Genetics, Maternal and Child Health (DiNOGMI), University of Genoa, Genoa, Italy; IRCCS Ospedale Policlinico San Martino, Genoa, Italy; Neuroradiology Unit, Fondazione IRCCS Ca’ Granda, Ospedale Maggiore Policlinico, Milan, Italy; Department of Pathophysiology and Transplantation, University of Milan, Milan, Italy; Department of Neurosciences “Rita Levi Montalcini”, University of Turin, Turin, Italy; Brain Research Imaging Centre, Cardiff University, Cardiff, UK

**Keywords:** glymphatic system, DTI-ALPS, quantitative susceptibility mapping, NODDI, Alzheimer’s disease

## Abstract

1.

**Background:** The glymphatic system facilitates clearance of metabolic waste and pathological proteins from the brain, and its dysfunction has been implicated in neurodegenerative disease. The diffusion tensor imaging–analysis along the perivascular space (DTI-ALPS) has been proposed as a non-invasive MRI marker of glymphatic flow, although its biological specificity remains uncertain. This study aimed to identify the determinants of DTI-ALPS and evaluate whether it primarily reflects white-matter (WM) microstructure rather than glymphatic flow in the context of neurodegeneration.

**Methods:** We examined 100 individuals referred to the Memory Clinic of the Policlinico Hospital in Milan for suspected dementia. All participants underwent a 3T-MRI protocol including 3D-T1-weighted and 3D-FLAIR imaging, double-shell diffusion-weighted imaging (b=1000/2000 s/mm²), and multi-echo gradient-echo sequences for quantitative susceptibility mapping. Within standard DTI-ALPS ROIs, we extracted DTI-ALPS values together with fractional anisotropy (FA), mean diffusivity (MD), and mode of anisotropy (MA) at both b-values, as well as neurite orientation and density imaging (NODDI) metrics, particularly the orientation dispersion index (ODI). WM microstructure was further characterized using the T1/FLAIR ratio and diamagnetic component of susceptibility (DCS).

**Results and conclusions:** DTI-ALPS correlated inversely with MA (r = –0.84 at b = 1000; r = –0.86 at b = 2000) and positively with ODI (r = 0.73). Moderate correlations with the T1/FLAIR ratio and DCS supported sensitivity to WM alterations. Factor analysis indicated that DTI-ALPS clustered with MA and ODI rather than forming a distinct factor, suggesting that DTI-ALPS primarily reflects WM diffusion dispersion and heterogeneity rather than glymphatic flow in the context of neurodegeneration.

**Keypoints:**

1. DTI-ALPS links strongly with mode of anisotropy at high b-values and orientation dispersion, indicating crossing-fibers loss
2. DTI-ALPS is moderately associated to T1/FLAIR ratio and diamagnetic susceptibility values, reflecting microstructural WM changes
3. DTI-ALPS reflects primarily WM features and may lack specificity for glymphatic flow in the context of neurodegeneration.

## 2. INTRODUCTION

The glymphatic system (GS) is a brain-wide perivascular network that mediates the flow and clearance of cerebrospinal fluid (CSF) and solutes through the brain parenchyma, facilitated by aquaporin-4 channels on astrocytic endfeet [1–3].

Its description in 2012[4] reshaped our understanding of cerebral fluid dynamics and was later complemented by the (re)discovery of meningeal lymphatic vessels in the dura[5,6], as well as the identification of direct connections linking the subarachnoid space with dural perisinus compartments[7], and the latter with the bone marrow within the thecal bone[8–10].

Together, these findings established the glymphatic–lymphatic model, with known roles in metabolic waste clearance[11] and immune surveillance of the brain[3,12]. Dysfunction of this system has been implicated in neurodegeneration, both through impaired clearance of pathological proteins such as amyloid-β[4] and tau[13,14], and through altered immune responses to neuropathology[12], underscoring its potential as a target for pharmacological intervention. Consequently, the in vivo visualization and assessment of glymphatic flow in humans have become a prominent focus of research.

Although the expression *study of the glymphatic system* has often been applied to approaches aimed at broadly characterizing CSF dynamics—from its production in the choroid plexus, to its movement along major subarachnoid vessels, and its drainage into meningeal lymphatic vessels—the core of the GS lies in the parenchymal passage of fluid, which has also proven to be the most elusive to measure.

Contrast-enhanced magnetic resonance imaging (CE-MRI) following intrathecal gadolinium administration is regarded as the gold standard for in vivo evaluation of the GS[15], as it enables quantification of CSF flow within the brain parenchyma under both physiological[16] and pathological conditions[17]. However, this technique is invasive, carries a risk of adverse effects and gadolinium-related toxicity, and requires prolonged serial MRI acquisitions over several hours or even days.

CE-MRI after intravenous gadolinium administration has been proposed as a less invasive alternative[18]. However, this approach remains poorly standardized[19], and its interpretation is complicated by the fact that contrast leakage from the vasculature into the central nervous system can occur through multiple pathways with distinct concentration–time profiles[20]. Moreover, the high intravascular gadolinium signal often obscures subtler extravascular changes, further complicating image analysis[20].

As a result, recent studies have increasingly focused on finding non-invasive biomarkers of GS function. Among these, the diffusion tensor imaging–analysis along the perivascular space (DTI-ALPS)[21] has gained popularity, mainly because of its technical simplicity. This method was developed to measure water diffusion along the perivascular spaces of medullary veins while reducing the confounding effect of white matter (WM) fiber orientation. Since its introduction in 2017, DTI-ALPS has been widely used, especially in research on neurodegenerative diseases[22–30]. In these studies, lower DTI-ALPS values have been linked cross-sectionally to poorer cognitive performance and greater disability, and longitudinally to faster clinical decline over time, particularly in Alzheimer’s disease (AD). These findings have been interpreted as supporting evidence that glymphatic dysfunction contributes to the pathogenesis of neurodegenerative diseases.

More recently, however, the actual sensitivity of this metric to glymphatic flow has been questioned[31,32]. Emerging data suggest that the DTI-ALPS index may be influenced by WM microstructural properties and damage[33–35], also considering its derivation from an anatomically complex region with numerous crossing fibers[36,37]. This may be particularly relevant in the context of neurodegenerative dementias. Although traditionally viewed as disorders of the grey matter, these conditions are characterize by early and prominent WM degeneration and demyelination[38,39]. In AD, for instance, WM damage is widespread[40] and appears to follow a “retrogenesis” model, in which degenerative processes mirror the reverse sequence of developmental myelination[41,42].

Diffusion-weighted imaging (DWI) is arguably the most sensitive technique to WM microstructural alteration. Most commonly, diffusion-related parameters are derived using the diffusion tensor imaging (DTI) model. Among DTI–derived metrics, fractional anisotropy (FA) and mean diffusivity (MD) have been widely used to study WM changes in neurodegenerative diseases, typically showing widespread decrease of FA and MD compatible with degeneration of WM tracts[40]. The mode of anisotropy (MA), a less commonly used DTI metric, complements FA by distinguishing between linear and planar diffusion patterns[43]. Whereas FA reflects the overall degree of diffusion anisotropy, MA characterizes the tensor shape and provides insight into local fiber geometry, particularly in regions of complex architecture[44].

DTI is a widely acknowledged model that, however, suffers from several limitations[45], particularly the partial volume effect of extracellular free water and the bias introduced by fiber dispersion. The Neurite Orientation Dispersion and Density Imaging (NODDI)[46] extends conventional DTI by modeling the diffusion signal into three biologically meaningful compartments: (i) the intra-neurite compartment, characterized by restricted diffusion within neurites; (ii) the extra-neurite compartment, reflecting hindered diffusion in the surrounding space; and (iii) the isotropic compartment, representing free water diffusion. Its key parameters—the neurite density index (NDI) and the orientation dispersion index (ODI)—quantify axonal density and angular variability of neurite orientation, respectively. Together, NDI and ODI provide more specific and interpretable insights into microstructural alterations, as they disentangle changes in neurite density and orientation dispersion that can produce similar effects on FA[47].

Structural MRI has been shown to provide complimentary information to DWI imaging in the assessment of WM degeneration[48], with the advantage of ease of accessibility. In particular, two imaging indices originally developed in the context of neuroinflammatory and demyelinating diseases have been proposed as structural markers of WM damage with possible applications in neurodegenerative diseases. The first is the ratio of T1- to T2-weighted[49] or fluid-attenuated inversion recovery (FLAIR) signal intensity[50,51], which has been shown to correlate with WM degeneration and poorer cognitive performance in AD and other neurodegenerative conditions[52,53].

The second is quantitative susceptibility mapping (QSM), an MRI technique that non-invasively quantifies tissue magnetic susceptibility, primarily reflecting contributions from iron and myelin[54]. However, conventional QSM does not resolve sub-voxel variations in susceptibility, and opposing diamagnetic (myelin-related) and paramagnetic (iron-related) sources may partially cancel each other, leading to artificially low or near-zero total values. To overcome this limitation, advanced algorithms have been developed to separate the diamagnetic (DCS) and paramagnetic (PCS) components of the QSM signal, enabling more specific assessment of myelin and iron content. This approach has shown promise as a potential tool for tracking pathological neurodegeneration across different brain regions and tissue types[55,56].

Against this background, in this study we sought to clarify the biological specificity of the DTI-ALPS index by assessing the potential confounding influence of WM microstructural alterations in the context of neurodegenerative disease. We examined a cohort of patients with suspected dementia, enriched for AD cases, and evaluated associations between DTI-ALPS and complementary MRI markers of WM damage within the same regions of interest (ROIs) used for DTI-ALPS computation. Specifically, we first examined correlations with DTI metrics at b=1000 and b=2000 to assess how their relationship with DTI-ALPS changes at higher diffusion weightings, which progressively suppress the contribution of freely diffusing water[57]. Then, we looked for a better explanation of these results with NODDI-derived indices reflecting neurite density, orientation dispersion and extracellular free water. Lastly, we compared DTI-ALPS with structural MRI markers of WM damage. This multimodal approach aimed to disentangle whether the DTI-ALPS index primarily reflects WM microstructural damage rather than glymphatic flow.

## 3. MATERIALS AND METHODS

### 3.1 Study participants

A total of 100 participants were recruited from individuals referred to the Neurodegenerative Diseases Unit of the Fondazione IRCCS Ca’ Granda Ospedale Maggiore Policlinico in Milan (Italy) for suspected dementia.

To be eligible for inclusion, subjects were required to have undergone the same MRI protocol on a 3T Philips Achieva dStream scanner (Philips Healthcare, Best, The Netherlands). The MRI protocol included:

1. a volumetric high-resolution T1-weighted Spoiled Gradient Echo (T1-SPGR);
2. a volumetric FLAIR;
3. a double-shell (b = 1000 and b = 2000) DWI (single-shot spin-echo echo-planar imaging) sequence acquired with opposite phase-encoding directions (posterior–anterior and anterior–posterior); and
4. a 3D multi-echo gradient echo (GRE) sequence suitable for QSM. Detailed MRI acquisition parameters are provided in **Table 1**.

**Table 1.** MRI protocol.

| Sequence / Protocol | 3D-T1 | 3D-FLAIR | DWI (EPI) | QSM (multi-echo GRE) |
| --- | --- | --- | --- | --- |
| Plane | Sagittal | Sagittal | Axial | Axial |
| TR (ms) | 8.2 | 5000 | 8400 | 40 |
| TE (ms) | 3.8 | 300 | 85 | 5.4-36.6 |
| $\Delta TE$ (ms) | - | - | - | 5.2 |
| Echoes | 1 | 1 | 1 | 7 |
| TI (ms) | - | 1700 | - | - |
| Slice thickness (mm) | 1 | 1 | 2.5 | 1 |
| Slice spacing (mm) | 1 | 1 | 0 | 0 |
| Flip angle ($^\circ$) | 8 | 120 | 90 | 18 |
| Matrix | 240x240 | 240x240 | 96x96 | 224x224 |
| b-values | - | - | 1000/2000 | - |
| Encoding directions | - | - | 32/32 | - |
| Phase encoding (main) | - | - | PA | - |

**Abbreviations:** FLAIR: fluid-attenuated inversion recovery; DWI: diffusion-weighted imaging; EPI: echo-planar imaging; QSM: quantitative susceptibility mapping; GRE: gradient echo; TR: repetition time; TE: echo time; TI: inversion time; PA: postero-anterior

At baseline, all participants underwent a standardized diagnostic assessment, including detailed medical history, general and neurological examination, comprehensive neuropsychological testing and mini-mental state examination (MMSE), MRI, and—when clinically indicated—CSF analysis and/or ^18^F-fluorodeoxyglucose positron emission tomography (FDG-PET) and/or amyloid-PET.

Participants underwent clinical follow-up every six months for at least 12 months. Based on longitudinal evaluation, individuals were classified as having a progressive neurodegenerative disorder (either of AD[58,59] or non-AD type[60–64]) or a non-degenerative condition, defined as stable mild cognitive impairment (sMCI), when no evidence of disease progression or longitudinal cognitive decline was observed.

Cognitively unimpaired (CU) participants presented with subjective cognitive complaints but exhibited no measurable impairment at formal neuropsychological testing nor evidence of an underlying neurodegenerative process after an extensive diagnostic work-up.

Exclusion criteria included: (i) MRI evidence of significant cerebrovascular pathology, cerebral amyloid angiopathy, or intracranial mass lesions; (ii) diagnosis of normal pressure hydrocephalus; and (iii) presence of severe motion artifacts compromising image quality.

### 3.2. DWI preprocessing, DTI metrics extraction and DTI-ALPS analysis

The DTI-ALPS index was automatically computed using the FSL-based pipeline developed by Liu et al[65].

Briefly, DWI were preprocessed in FSL[66,67]. After DICOM-to-NIfTI conversion, images were corrected for susceptibility-induced field inhomogeneities and subject motion using topup[68] and eddy[69]. A brain mask was then generated from the averaged b0 volumes using BET[70].

Diffusion tensor fitting was performed with dtifit considering the b=1000 and b=2000 s/mm² shells separately. This produced voxel-wise maps of FA, MD, and full diffusion tensors for each b-value.

FA maps were normalized to the high-resolution FMRIB58_FA standard space image using linear and non-linear registrations (FLIRT and FNIRT)[71], and tensors were transformed into the same space with vector correction. Directional tensor components (Dxx, Dyy, Dzz) were extracted, and all indices were computed within default ROIs corresponding to 2.5-mm-radius spheres located in projection and association fibers of both hemispheres, as defined in the Johns Hopkins University (JHU) atlas.

The DTI-ALPS index was calculated separately for each b-value (referred to as DTI-ALPS_1000_ and DTI-ALPS_2000_ throughout the text) as the ratio between diffusivity along the x-axis and the mean diffusivity along the orthogonal (y and z) axes in the 4 ROIs, yielding a global value. Mean FA and MD were also computed across the four ROIs for both b-values (FA_1000_, FA_2000_, MD_1000_, and MD_2000_).

Additionally, MA was calculated following the formulation described by Ennis and Kindlmann to obtain mean global MA across the four ROIs for both diffusion weightings (MA_1000_ and MA_2000_)[43]. Beyond the magnitude of anisotropy captured by FA, the MA describes the tensor’s geometric configuration, varying from linear (MA ≈ +1) to planar (MA ≈ –1) and isotropic (MA ≈ 0), and reflects the relative distribution of the diffusion eigenvalues.

### 3.3 T1 and FLAIR images preprocessing and T1/FLAIR ratio calculation

T1/FLAIR ratio maps were processed following a multistep pipeline available at https://github.com/treanus/KUL_NIS, as referenced in Cappelle[51]. Briefly, intensity nonuniformities due to field inhomogeneity were corrected using the N4 bias field correction algorithm implemented in ANTs[72]. The bias-corrected FLAIR images were then rigidly coregistered to the corresponding T1 volumes using antsRegistration.

T1 images were subsequently nonlinearly normalized to the MNI152 (ICBM) standard space, producing forward and inverse deformation fields.

To improve inter-subject comparability, intensity calibration was carried out within brain masks of both the subject and the MNI template, as implemented in the calib-nonlin3 procedure of the original code. During this step, the intensity histogram of each subject was nonlinearly adjusted to match that of the reference template, in order to standardize intensity distribution across subjects and minimize variations related to receiver gain or acquisition conditions. Calibrated T1 and FLAIR images were then combined voxel-wise to compute T1/FLAIR ratio maps. These ratio maps were spatially normalized to MNI space using the forward deformation fields derived from T1 normalization, and the mean T1/FLAIR ratio was extracted within the four DTI-ALPS ROIs (referred to as T1/FLAIR_ratio_).

### 3.4 DCS map calculation

DCS maps were derived using the DECOMPOSE algorithm developed by Chen et al.[73]. In this method, QSM reconstructed at each echo is used to fit a three-pool signal model representing paramagnetic (primarily iron-related), diamagnetic (primarily myelin-related), and magnetically neutral tissue compartments. Through this framework, the relative contributions of each magnetic source to the total susceptibility can be estimated voxel-wise.

For the DECOMPOSE reconstruction, only the first five echoes were used, as later echoes typically exhibited lower signal-to-noise ratio and greater phase instability[55]. DCS values were kept in their native units without normalization to a reference region, avoiding assumptions about brain areas presumed to be spared from neurodegenerative changes[54].

The resulting DCS maps were then coregistered to the corresponding native T1-weighted images and normalized to the MNI152 (ICBM) standard space with the same modalities applied for the T1/FLAIR_ratio_ calculation. All maps were visually inspected to confirm image quality, artifact suppression, and spatial correspondence prior to inclusion in subsequent analyses.

Subsequently, a histogram-based characterization of DCS values was performed. For each participant, the mean (DCS_mean_), median (DCS_med_), standard deviation (DCS_std_), kurtosis (DCS_kurt_), skewness (DCS_skew_), and the 10th (DCS_p10_), 25th (DCS_p25_), 75th (DCS_p75_), and 90th (DCS_p90_) percentiles were extracted by averaging values within the four DTI-ALPS ROIs[74].

### 3.5 NODDI metrics extraction

Microstructural diffusion parameters were estimated using the NODDI model[46] implemented within the AMICO (Accelerated Microstructure Imaging via Convex Optimization) framework[75]. The AMICO pipeline was applied to DWI that had been preprocessed for eddy current, motion, and susceptibility-induced distortion correction.

Within the AMICO framework, the NODDI model was fitted assuming three diffusion compartments: (i) an intra-neurite compartment representing restricted diffusion within neurites, (ii) an extra-neurite compartment modeling hindered diffusion in the surrounding tissue, and (iii) an isotropic compartment reflecting free water diffusion.

For each subject, the following parametric maps were generated: intracellular volume fraction, reflecting neurite density (NDI); orientation dispersion (ODI), quantifying neurite orientation variability; and isotropic volume fraction (F_ISO_), representing free water content. NODDI maps were normalized to the high-resolution FMRIB58_FA standard space image using the inverse DTI transformation matrix, ensuring alignment with other diffusion-derived metrics. Average NDI, ODI, and F_ISO_ values were then extracted across the four DTI-ALPS ROIs.

### 3.6 Statistical analysis

All statistical analyses were performed using R Studio (version 2025.09.1+401, R Foundation for Statistical Computing, Vienna, Austria). Data were first examined using descriptive statistics and visual inspection of distributions to assess normality.

Exploratory analyses were conducted to evaluate associations between the main study variables and demographic factors. Associations with age were tested using linear models (including both linear and quadratic terms to capture potential nonlinear effects), and associations with sex were assessed using two-sample t-tests. Pairwise correlations among continuous variables were computed using Pearson’s correlation coefficients, with two-tailed p-values adjusted for multiple comparisons using the False Discovery Rate (FDR) correction. Statistical significance was set at p < 0.05 (FDR-corrected).

Variables that showed significant correlations with DTI-ALPS_1000_ after FDR correction were further analyzed using multivariate linear regression models after scaling, with the DTI-ALPS_1000_ as the primary dependent variable, adjusting for age (linear and quadratic terms) and sex. Model performance was assessed by computing the adjusted coefficient of determination (adjR²), Akaike information criterion (AIC), and Bayesian information criterion (BIC).

An exploratory factor analysis (EFA) was then performed on the residuals of variables that remained significant after the regression analyses, thus using data adjusted for the effects of age and sex. We note that this represents a more aggressive correction, owing to potential shared variance between WM metrics and age. The suitability of the dataset for factor analysis was verified using the Kaiser–Meyer–Olkin (KMO) measure of sampling adequacy (threshold > 0.6, both overall and for each variable) and by confirming the absence of multicollinearity (pairwise r <0.8). The EFA was conducted with the psych package in R[76]. The optimal number of factors to retain was determined using the fa.parallel function based on parallel analysis. Factor extraction was performed with the minimum residual (minres) method, and factors were rotated using oblimin rotation to allow for correlated components. Model fit was assessed using Root Mean Square Error of Approximation (RMSEA), Tucker–Lewis Index (TLI) and Root Mean Square Residual (RMSR).

## 4. RESULTS

### 4.1 Demographic and clinical characteristics of the study population

Of the 100 subjects included in the study, 20 were CU elderly subjects, 21 sMCI patients, 39 AD patients, 20 patients with non-AD neurodegenerative conditions. The demographics for the whole cohort and each group separately are detailed in **Table 2**.

**Table 2.** Clinical and demographical data of the patients.

|  | All<br>n=100 | CU<br>n=20 | sMCI<br>n=21 | AD<br>n=39 | Non-AD<br>n=20 | Statistic<sup>a</sup> |
| --- | --- | --- | --- | --- | --- | --- |
| Age, years (Q1,Q3) | 73 (66,76) | 69 (60,75) | 74 (69,76) | 62 (66.5, 76.5) | 75.5 (70.75,78) | $\chi^2$ 6.43<br>$p$ 0.09 |
| Sex, Females:Males | 54:46 | 7:13 | 12:9 | 22:17 | 13:7 | $\chi^2$ 4.01<br>$p$ 0.26 |
| Years of Education (Q1,Q3) | 13 (8,18) | 13 (8,18) | 8 (8,13) | 13 (12,18) | 13 (7.75,14.25) | $\chi^2$ 6.82<br>$p$ 0.08 |
| MMSE at MRI (Q1,Q3) | 26* (23,28) | 29 (28.75,30) | 26 (25,28) | 23 (21, 26) | 24.3 (22,26) | $\chi^2$ 40.72<br>$p$ <0.001 |
| FDG-PET, Y:N | 46:54 | 8:12 | 9:12 | 15:24 | 14:6 | - - |
| FBB-PET, Y:N | 27:73 | 7:13 | 5:16 | 11:28 | 4:16 | - - |
| CSF, Y:N | 48:52 | 5:15 | 8:13 | 24:15 | 11:9 | - - |

**Abbreviations:** *CU*: cognitively unimpaired; *sMCI*: stable mild cognitive impairment; *AD*: Alzheimer's disease; *Non-AD*: Non-Alzheimer's disease dementias; *MMSE*: mini-mental state examination; *FDG-PET*: fluorodeoxyglucose positron emission tomography; *FBB-PET*: fluorbetaben positron emission tomography; *CSF*: cerebrospinal fluid. *a*: Reported P-values from one-way between-groups ANOVA for age, years of education, MMSE at MRI, and chi-square test for independence for sex, FDG-PET, FBB-PET and CSF.

There was no significant difference in sex distribution, age and years of education across the groups. A significant difference in MMSE at the time of MRI was observed (p<0.001), with pairwise comparisons revealing that the CU group had significant higher scores than all the other groups (p<0.001 for all comparisons) and that sMCI patients had higher values compared to AD patients (p<0.014).

All DTI measures at both b-values showed a linear effect of age (**see Supplementary figure 1 and 2** for R^2^ for linear and quadratic components), except for FA. ODI and NDI, but not F_ISO_, showed a linear reduction with age.

Among structural metrics, T1/FLAIR_ratio_ tended to decrease and DCS_kurt_ to increase quadratically with age, while DCS_std_ show a linear increase and DCS_skew_ a linear decrease with age. All other DCS metrics showed no relation with age.

Many measures exhibited a main effect of sex. Particularly, FA_1000_, FA_2000_, MA_1000_, were higher in men, while DCS_mean_, DCS_med_, DCS_p10_, DCS_p25_, DCS_p75_, DCS_p90_ and T1/FLAIR_ratio_ were higher in females. There were no sex differences in NODDI related metrics.

### 4.2 DTI-ALPS_1000_ is inversely associated with FA and MD in the DTI-ALPS ROI

**Figure 1** shows the correlations between the main study variables, and **Figure 2** displays the strongest associations through size- and symbol-coded scatterplots. A reduction in the DTI-ALPS_1000_ values was associated with an increase in FA_1000_ and MD_1000_ (r=-0.40; r=-0.56 respectively, all p_FDR_<0.001). Stronger correlations were found with FA_2000_ and MD_2000_ (r=-0.53; r=-0.60 respectively, all p_FDR_<0.001). After correction for age and sex in multivariate models, FA and MD at both b-values remained significant predictors of DTI-ALPS_1000_. (**Supplementary table 1**). AIC and BIC were consistently lower and adjR^2^ higher when considering measures extracted at b=2000.

**Figure 1.**
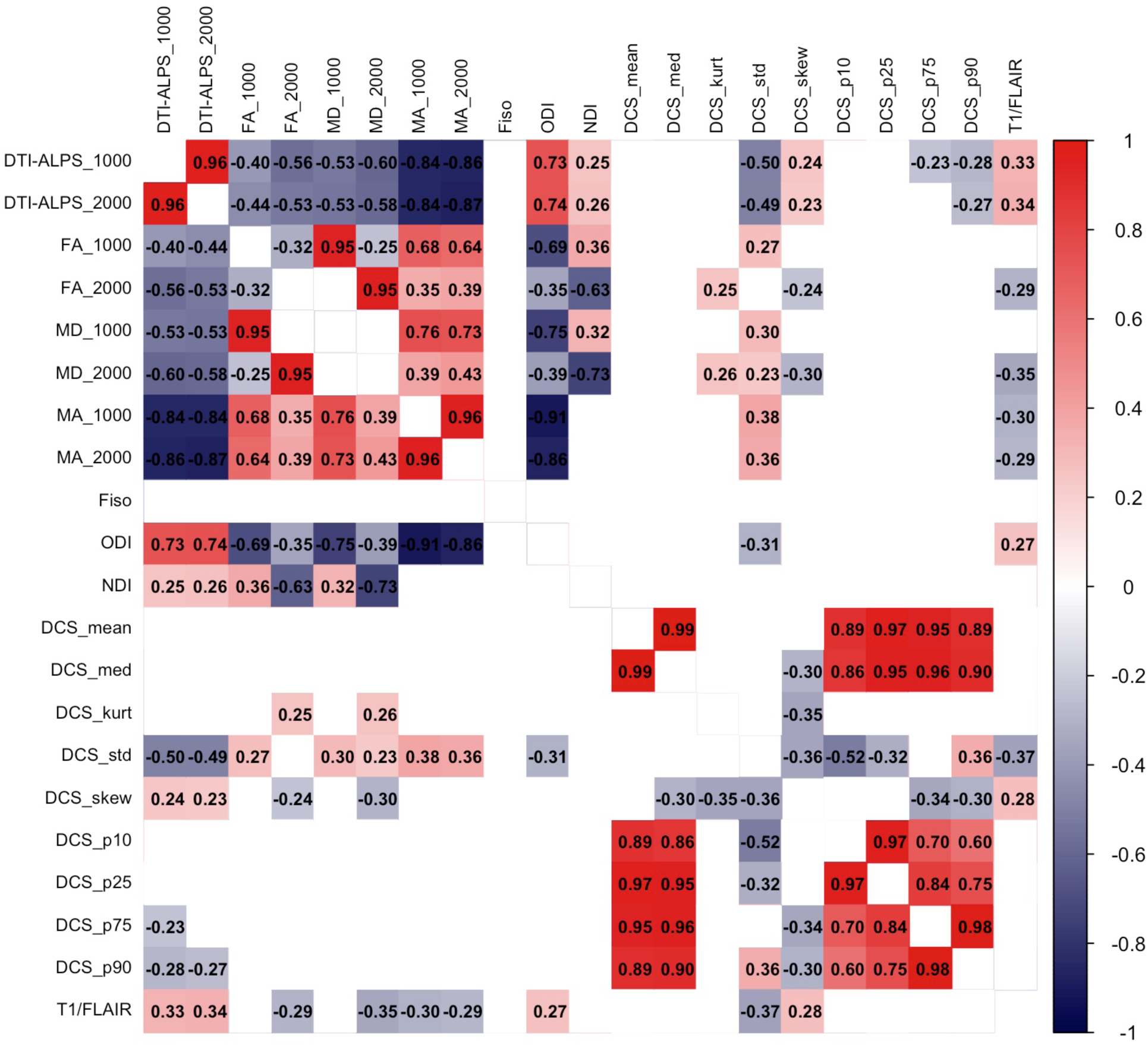
Correlation matrix of variables of interest. Positive correlations are shown in red and negative correlations in blue. Only statistically significant associations after FDR correction are displayed. Pearson’s r values are reported within the squares.

**Figure 2.**
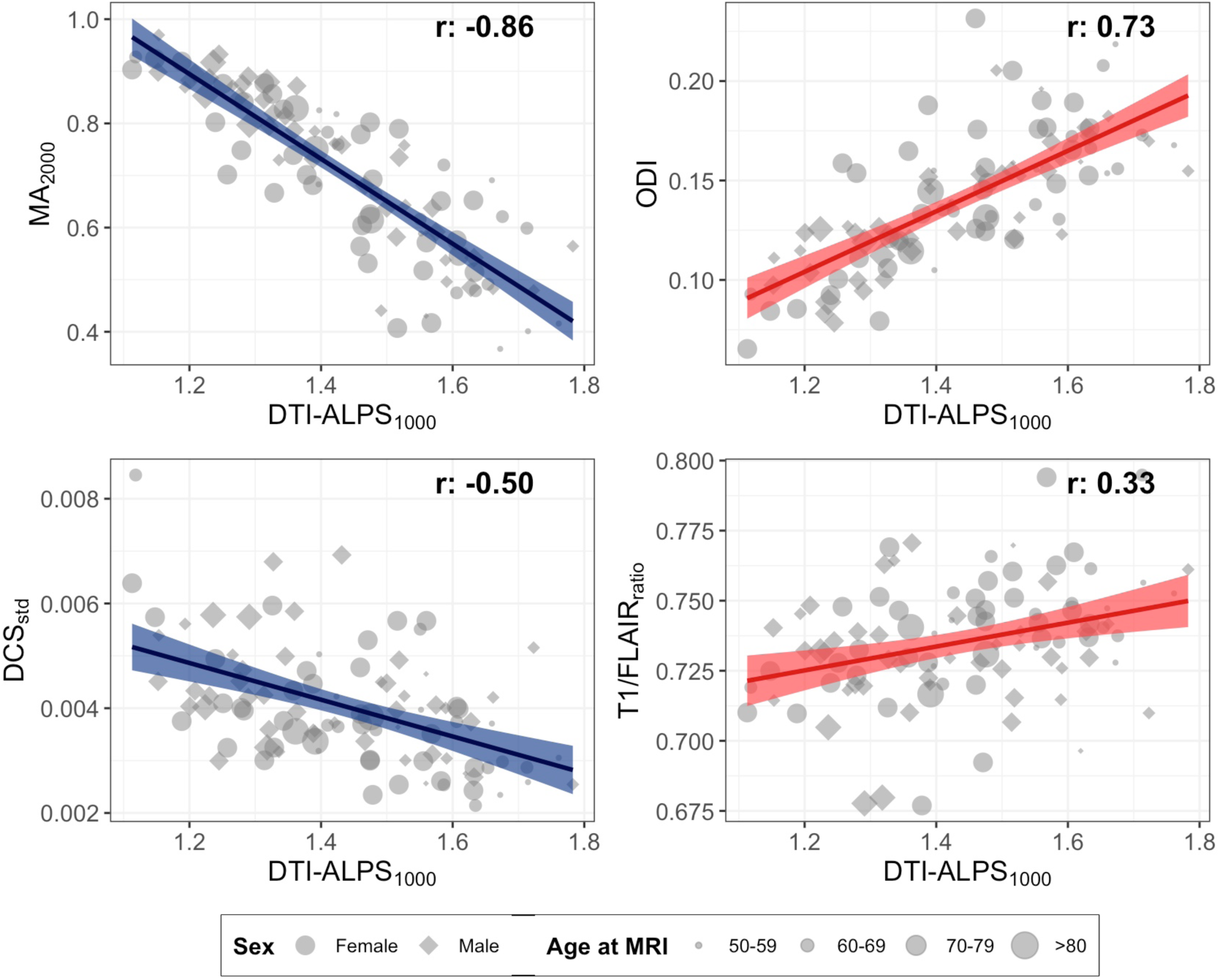
Bubble plots illustrating relationships between DTI-ALPS_1000_ and selected variables. Size scales with age, and marker shape denotes sex. Negative and positive correlations with DTI-ALPS_1000_ are shown by blue and red regression lines, respectively, with shaded areas representing standard error.

### 4.3 DTI-ALPS_1000_ is strongly and inversely associated with MA in DTI-ALPS ROIs

A reduction in DTI-ALPS_1000_ values was strongly associated with an increase in MA_1000_ and even higher increases in MA_2000_ (r = –0.84 and r = –0.86, respectively; all p_FDR_<0.001). Both MA_1000_ and MA_2000_ remained robust independent predictors of ALPS after adjustment for age and sex (**Supplementary table 1**). Model fit indices were consistently lower, and explained variance higher, when measures derived from the b=2000 shell were used.

### 4.4 DTI-ALPS_1000_ is highly and directly associated with ODI in the ALPS ROIs

DTI-ALPS_1000_ showed a strong positive correlation with ODI (r 0.73, p_FDR_<0.001). We also found a weaker positive association with NDI (r 0.25, p_FDR_=0.030). ODI showed also a strong negative correlation with MA at both b=1000 and b=2000 (r -0.91 and -0.86 respectively, all p_FDR_<0.001).

In the multivariate model, only ODI remained a significant independent predictor of DTI-ALPS_1000_ (**Supplementary table 1**).

### 4.5 DTI-ALPS_1000_ values show moderate associations with structural WM markers

When considering structural MRI metrics, DTI-ALPS_1000_ values showed a direct association with T1/FLAIR_ratio_ (r=0.33, p_FDR_=0.003) and with DCS_skew_ (r=0.24, p_FDR_=0.037). On the other hand, DTI-ALPS_1000_ showed a strong negative correlation with DCS_std_ (r=-0.50, p_FDR_<0.001) and moderate association with DCS_p75_ and DCS_p90_ (r=-0.23, p_FDR_=0.045; r=-0.28, p_FDR_=0.012 respectively).

T1/FLAIR_ratio_ was negatively associated with MA_1000_ (r -0.30, p_FDR_=0.006 FDR), MA_2000_ (r - 0.29, p_FDR_=0.010), MD_1000_ (r -0.29, p_FDR_=0.008) and MD_2000_ (r -0.35, p_FDR_=0.001), while positively correlated with ODI (r 0.27, p_FDR_=0.018). DCS_std_ showed the opposite pattern: it was positively associated with MA_1000_ (r 0.38, p_FDR_<0.001), MA_2000_ (r 0.36, p_FDR_<0.001), FA_1000_ (r 0.27, p_FDR_ =0.016), FA_2000_ (r 0.30, p_FDR_=0.008) and MD_2000_ (r 0.23, p_FDR_=0.046), while negatively correlated with ODI (r -0.31, p_FDR_=0.006)

DCS_std_, DCS_p75_, DCS_p90_ and T1/FLAIR_ratio_ remained significant predictors of DTI-ALPS_1000_ after correcting for age and sex. By contrast DCS_skew_ became unsignificant (**Supplementary table 1**).

### 4.6 DTI-ALPS_1000_ associates with diffusion anisotropy metrics in factor analysis

Prior to factor extraction, three variables were excluded: FA and MD at both b-values, which showed low sampling adequacy (KMO < 0.6) and DCS_p75_, which exhibited high collinearity with DCS_p90_. ODI, MA_1000_ and MA_2000_ showed high collinearity, leading to Heywood case when kept together in the model. However, as these were the primary variables of interest, rather than excluding one, we conducted two separate EFAs—one retaining ODI and the other retaining MA_2000_ (MA_1000_ was excluded due to high collinearity and its weaker correlation with DTI-ALPS_1000_).

Considering ODI, a two-factor solution provided the best fit to the data (χ² = 0.42, p = 0.52; RMSEA = 0.00, 90% CI = [0, 0.23]; TLI = 1.06; RMSR = 0.01), accounting for 54% of the total variance (Factor 1 = 29%; Factor 2 = 25%). Factor 1 loaded primarily on DTI-ALPS_1000_ (β=0.62) and ODI (β=1.01), whereas Factor 2 was driven by DCS_std_ (β=0.90) and DCS_p90_ (β=0.42), with smaller contributions from T1/FLAIR_ratio_ (β=–0.33). The two factors were moderately correlated (r –0.32), indicating partially overlapping latent constructs. The model demonstrated excellent fit and high factor score reliability (minimum correlation = 0.66–0.97).

When MA_2000_ was retained, results were almost alike. A two-factor solution provided an adequate fit to the data (χ² = 2.6, p = 0.11; RMSEA = 0.13, 90% CI = [0, 0.33]; TLI = 0.90; RMSR = 0.02), explaining 56% of the total variance (Factor 1 = 33%; Factor 2 = 23%). Factor 1 loaded primarily on MA_2000_ (β=1.03) and inversely on DTI-ALPS_1000_ (β=–0.75), while Factor 2 was driven by DCS_std_ (β=0.83) and, to a lesser extent, by DCS_p90_ (β=0.48) and T1/FLAIR_ratio_ (β=–0.35). The two factors were moderately correlated (r 0.39), suggesting partially overlapping latent dimensions. Model fit indices indicated good adequacy, with a TLI of 0.895 and strong factor score reliability (minimum correlation = 0.53–0.99)(**Figure 3**).

**Figure 3.**
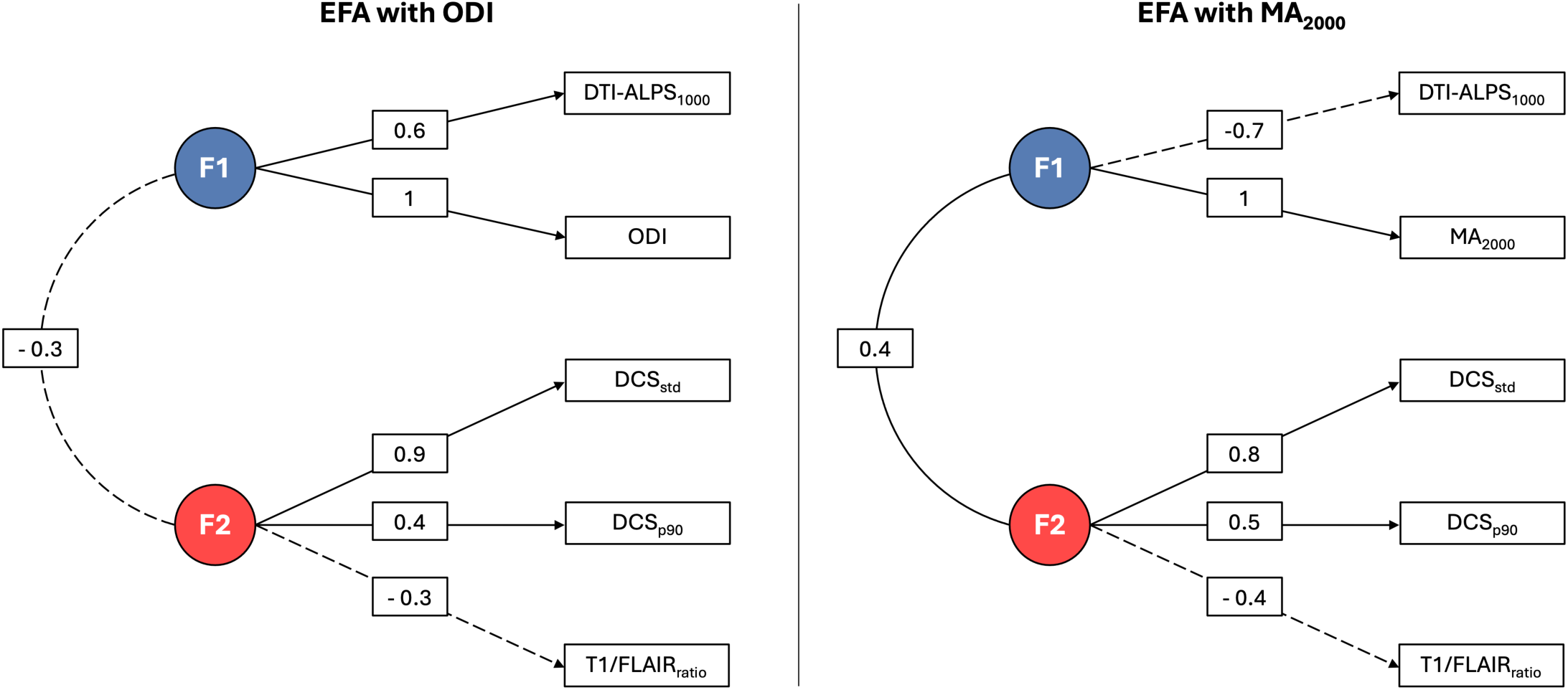
Exploratory factor analysis models including either ODI or MA_2000_. The two models show latent factors (F₁, F₂) and their standardized loadings onto diffusion and structural metrics. Dashed arrows indicate negative loadings, and numbers represent standardized factor loadings or inter-factor correlations. Model including ODI (left); model including MA_2000_ (right).

## 5. DISCUSSION

In this study, we aimed to characterize the determinants of DTI-ALPS changes in neurodegenerative diseases using a clinical MRI protocol incorporating multi-shell DWI and structural MRI. We found that DTI-ALPS, calculated at b=1000, is highly associated with DTI measure of diffusion dispersion, particularly MA and ODI, and demonstrates a moderate relationship with structural markers of WM microstructural damage, namely T1/FLAIR_ratio_ and DCS.

Overall, our findings suggest that DTI-ALPS predominantly reflects loss of crossing fibers and WM microstructural damage within the ROIs where it is calculated, rather than serving as a direct measure of glymphatic function, as originally proposed. We substantiate this interpretation in greater detail in the following paragraphs.

In the original paper by Taoka, DTI-ALPS was proposed as a marker of glymphatic activity based on the hypothesis that in an area “in which conformation of the perivascular space, represented by medullary veins on SWI, and the major white matter tracts were perpendicular to each other”, the “ratio of x-axis diffusivity in the projection fibers (i.e. corticospinal tract) and association fibers (i.e. superior longitudinal fasciculus) area (Dxproj and Dxassoc) to the diffusivity which is perpendicular to them (Dyproj and Dzassoc) would express the influence of the water diffusion along the perivascular space which will reflect activity of the glymphatic system in the individual cases”[21].

Therefore, for DTI-ALPS to truly represent glymphatic activity, changes in the index should primarily arise from variations in x-axis diffusivity (Dxx), which is hypothesized to be most sensitive to perivascular fluid movement.

Considering the main orientation of eigenvector-2 in these tracts, the DTI-ALPS index can be approximated as ≈ λ2/λ3, assuming similar λ2/λ3 ratios for the two fibers types[35].

Consequently, and all else being equal, a reduction in glymphatic flow would be expected to increase FA anisotropy within DTI-ALPS-ROIs and decrease MD, due to the reduction of λ2 component. In addition, it would result in a reduction of radial asymmetry (i.e the ratio between λ2 and λ3)[33,35].

Even if reduced glymphatic flow were instead to result in fluid stagnation and an increase in free water content, the ensuing diffusion pattern would be expected to become more isotropic and once again characterized by decreased FA and increased MD. This latter pattern is typical of WM damage and axonal degeneration, where expansion of the extracellular space occurs secondary to axonal loss[40].

In our cohort, however, we observed an inverse correlation between DTI-ALPS index and both FA and MD within the DTI-ALPS ROIs, a finding that may initially seem counterintuitive. Nonetheless, a small number of prior studies have reported parallel increases in FA[44,77–79] and MD[78] in AD, Parkinson’s disease and progressive supranuclear palsy[34], particularly in the centrum semiovale[78].

These parallel increases of FA and MD have been attributed to the selective degeneration of crossing fibers—a hypothesis supported by multiple converging lines of evidence: (i) the centrum semiovale, from which DTI-ALPS is derived, is densely packed with crossing bundles[36,37]; (ii) AD is characterized by early degeneration of intracortical association fibers, with relative preservation of corticofugal and corticopetal tracts[80] (iii) Increases in FA and MD were accompanied by a corresponding rise in MA[44,77]; (iv) quantitative tractography indicated that this effect was directly related to the selective sparing of motor-related projection fibers[77,78].

Indeed, the colocalized increase in MA within such crossing-fiber regions suggests a transition of the WM bundles toward a more linear (“cigar-like”) shape, consistent with loss of crossing fibers.

In line with this evidence, we found that MA alone explained more than 75% of the variance in DTI-ALPS, strongly suggesting that DTI-ALPS primarily reflects the integrity of crossing fibers rather than perivascular diffusion.

Given the assumptions outlined above and the sensitivity of DTI to extracellular free water, a reduction in λ₂ diffusivity—presumed to reflect perivascular glymphatic flow (Dxx)—could theoretically increase FA and thus MA. However, several lines of evidence argue against a substantial glymphatic contribution to the parallel changes observed in MA and DTI-ALPS. First, if DTI-ALPS were sensitive to perivascular diffusion, its correlations with DTI metrics would be strongest at lower b-values (i.e. b=1000 s/mm²). Instead, we found stronger associations at higher b-values (i.e. b=2000 s/mm²), reinforcing the view that DTI-ALPS is primarily driven by axonal microstructure, as already reported[33–35].

Second, perivascular spaces occupy only about 1% of the WM volume[31,81], representing a negligible fraction of the DTI-ALPS ROIs too. Even assuming some sensitivity of DTI-ALPS to glymphatic dynamics, the association with MA_2000_ should display considerable interindividual variability, given (i) expected differences in the relative contributions of glymphatic and microstructural effects across subjects, and (ii) opposing influences of glymphatic flow reduction on FA—potentially increasing FA through decreased λ₂ while simultaneously decreasing FA through extracellular fluid accumulation[25,26]. This would produce greater scatter in the DTI-ALPS–MA relationship than we observed.

Third, we found moderate correlations between DTI-ALPS and structural MRI indices of WM integrity, particularly T1/FLAIR_ratio_ and DCS_std_. Although the association between DTI-ALPS and conventional DTI metrics was weaker—likely reflecting the greater sensitivity of diffusion imaging to subtle microstructural changes[78]—these findings support a link between DTI-ALPS and WM damage. Specifically, the positive correlation between DTI-ALPS and the T1/FLAIR_ratio_ suggests that lower DTI-ALPS values correspond to reduced tissue integrity. The T1/FLAIR_ratio_ has been proposed as a practical alternative to the T1/T2 ratio for assessing myelin-related properties[51], given the widespread use of volumetric FLAIR over T2-weighted sequences in current MRI protocols. Despite concerns about its specificity for myelin, based on comparisons with other myelin mapping modalities[82–84] and post-mortem studies[85–87], the T1/FLAIR_ratio_ remains a valuable composite marker of tissue integrity reflecting contributions from myelin, iron, calcium, axonal density, and extracellular content[51,83]. It has also been proposed as a sensitive, clinically accessible marker of WM damage in neurodegenerative disorders[52,53]. Accordingly, its reduction in DTI-ALPS ROIs supports the interpretation that WM damage confounds DTI-ALPS metrics. A similar interpretation applies to DCS. We hypothesized that lower DTI-ALPS values would correspond to higher DCS—reflecting myelin loss, as previously reported in AD[56]. Although a trend toward increased mean and median DCS within DTI-ALPS ROIs was observed, it did not reach statistical significance. However, DTI-ALPS showed a strong inverse correlation with DCS_std_ and a moderate inverse correlation with DCS_p90_. Together, these findings suggest that (i) a reduction of DTI-ALPS is associated with a shift toward higher DCS values, consistent with demyelination[56,88], and (ii) lower DTI-ALPS coincides with greater DCS variability, indicative of increased microstructural heterogeneity, in agreement with T1/FLAIR_ratio_ results. While susceptibility measures can be affected by fiber orientation relative to the main magnetic field, averaging across two orthogonal ROI sets likely minimized this confound[89].

Fourth, DTI-ALPS showed a strong positive correlation with ODI, which was itself closely related to MA. Reduced ODI has been consistently reported in neurodegenerative diseases and interpreted as a consequence of selective loss of crossing fibers[47,90,91], leading to greater alignment of remaining axons. ODI has also demonstrated stronger histological validity than DTI metrics[92,93], likely because it corrects for the biasing effect of extracellular free water. Moreover, as ODI derives from the neurite compartment within the NODDI framework, its relationship with DTI-ALPS reinforces the view that ALPS primarily captures axonal geometry rather than perivascular flow.

Although MA, ODI, and structural indices of WM damage accounted for most of the DTI-ALPS variance, a small portion remained unexplained. This residual variance, however, should not automatically be attributed to glymphatic activity, as it may instead reflect unmodelled biological or methodological factors.

Indeed, EFA clearly revealed that DTI-ALPS do not form a distinct component but instead clusters with MA and ODI, indicating that it largely reflects neurite orientation and diffusion coherence rather than a separate physiological process.

Moreover, it is important to note that the DTI-ALPS index primarily reflects the radial asymmetry of underlying tracts[33,35], which depends not only on crossing fibers but also on microscopic features of the fascicles—such as dispersion anisotropy and axonal undulation. These microscopic features may be less effectively detected by MA and ODI, requiring more advanced modeling taking into account also non isotropic dispersion[94]. Finally, the small fraction of variance left unexplained by MA, ODI and structural MRI markers may simply reflect measurement noise. In this context, the imperfect alignment between WM tracts and reference axes merits specific consideration. Because diffusion tensor eigenvectors rarely coincide exactly with these axes, even minor misalignments can introduce discrepancies between MA and DTI-ALPS despite their shared microstructural basis. To verify this assumption, we calculated a modified version of MA substituting eigenvalues with corresponding diffusion axes in the ROIs, which led to an increase in the proportion of DTI-ALPS variance explained—particularly at b=2000—further supporting their common structural origin (**Supplementary table 2**).

## 6. LIMITATIONS

This study has several limitations. First, diffusion data were acquired with a maximum b-value of 2000 s/mm², which may not fully suppress the contribution of freely diffusing water. Higher b-values could enhance the sensitivity of diffusion metrics to tissue microstructure; however, the acquisition protocol was constrained by clinical MRI parameters to ensure feasibility in a patient population.

Second, the NODDI model used here assumes isotropic orientation dispersion, and thus does not account for potential anisotropic components that may better capture the complexity of white matter fiber geometry. Incorporating more advanced diffusion models (i.e Bingham NODDI)[94] could provide a more accurate description of microstructural dispersion patterns. Moreover, it could test if DTI-ALPS is able to capture reduction of dispersion linked to anisotropic components, as has been proposed.

Finally, the cross-sectional design limits the ability to determine how WM microstructural confounders influence the predictive validity of the DTI-ALPS index over time. Longitudinal studies are needed to determine whether the DTI-ALPS index maintains its predictive value in this context.

## 7. CONCLUSIONS

In conclusion, our study demonstrates that DTI-ALPS measures are strongly associated with WM microstructural integrity, as assessed through both DTI and structural MRI. Specifically, our findings suggest that DTI-ALPS metrics primarily reflect loss of crossing fibers and increased microstructural heterogeneity within DTI-ALPS ROIs, challenging its interpretation as a direct marker of glymphatic activity as originally proposed by Taoka. Nevertheless, our intention is not to fully dismiss ALPS—which remains the only putative glymphatic marker tested against intrathecal gadolinium[95]—but rather to emphasize the need for further validation and consensus. The sensitivity of the DTI-ALPS index to WM geometry (i.e., diffusion asymmetry) may still prove valuable for investigating WM damage in neurodegenerative diseases. Moreover, given the dependence of glymphatic clearance on normal microstructural anatomy, DTI-ALPS could still be proved a useful marker of *glymphatic dysfunction secondary to WM injury*, provided its correlation with gadolinium clearance in intrathecal CE-MRI is confirmed. However, current evidence—including our own—suggests that DTI-ALPS is influenced by WM microstructure and may not reliably represent a *direct marker of glymphatic flow*.

## DECLARATIONS

### Ethics approval and consent to participate

all procedures performed in studies involving human participants were in accordance with the ethical standards of the institutional and/or national research committee and with the 1964 Helsinki declaration and its later amendments or comparable ethical standards. Written informed consent was obtained from all participants or their legal proxies. The study was approved by the local Ethics Committee (Comitato Etico Area 2 Milano, approval N 859_2021, dated 14 September 2021).

### Availability of data and materials

the datasets used in this study are available from the corresponding author upon reasonable request.

## Funding

this research was supported by grants from the Italian Ministry of Health (Ricerca Corrente to AA and DG, Grant PNRR-POC-2023-12377354 to DG) and from the Fondazione Regionale per la Ricerca Biomedica (FRRB)-UNMET MEDICAL NEEDS—Regione Lombardia (Acronym: MAINSTREAM, Project ID: 3430931) to TC.

### Authors’ contributions

LS and AA conceived and designed the study. LS interpreted the data, performed the statistical analyses, and drafted the manuscript. GF and GBoc conducted neuroimaging data preprocessing and analysis. MC and GBof revised the content of the manuscript as field expert. TC, LG, MD, AP, and MP assisted with the acquisition of clinical data. ES and FT acquired the MRI data. AA, MB, and DG critically revised the manuscript for important intellectual content. All authors read and approved the final version of the manuscript.

## Acknowledgements

no acknowledgments to declare.

---

**Supplementary figure 1.** Scatterplot grid showing associations between diffusion-derived metrics and age. Each panel depicts the relationship between a diffusion metric and age, modeled with both linear and quadratic components. Solid lines represent fitted polynomial regression curves.

**Supplementary figure 2.** Scatterplot grid showing associations between structural MRI-derived metrics and age. Each panel depicts the relationship between a structural MRI metric and age, modeled with both linear and quadratic components. Solid lines represent fitted polynomial regression curves.

