## Supplementary tables and figures for "DTI-ALPS PRIMARILY REFLECTS WHITE MATTER DIFFUSION DISPERSION AND MICROSTRUCTURAL HETEROGENEITY IN NEURODEGENERATION: INSIGHTS FROM MULTI-MODAL MRI"

#### 10. SUPPLEMENTARY MATERIALS

**Supplementary table 1. Multivariate analysis results with DTI-ALPS$_{1000}$ as dependent variable**

**FA_1000**

| term | estimate | std.error | statistic | p.value sig |
|---|---|---|---|---|
| (Intercept) | -0.761 | 5.751 | -0.132 | 0.895 |
| FA_1000 | -0.348 | 0.084 | -4.134 | **<0.001*** ** |
| Age at MRI | 0.089 | 0.172 | 0.518 | 0.606 |
| Age at MRI^2 | -0.001 | 0.001 | -0.855 | 0.394 |
| Sex | -0.102 | 0.168 | -0.607 | 0.545 |
| | | **adjR2**: 0.341 | **AIC**: 248.974 | **BIC**: 264.605 |

**FA_2000**

| term | estimate | std.error | statistic | p.value sig |
|---|---|---|---|---|
| (Intercept) | -0.409 | 5.360 | -0.076 | 0.939 |
| FA_2000 | -0.463 | 0.079 | -5.834 | **<0.001*** ** |
| Age at MRI | 0.073 | 0.160 | 0.453 | 0.652 |
| Age at MRI^2 | -0.001 | 0.001 | -0.787 | 0.433 |
| Sex | -0.039 | 0.158 | -0.245 | 0.807 |
| | | **adjR2**: 0.427 | **AIC**: 234.898 | **BIC**: 250.529 |

**MD_1000**

| term | estimate | std.error | statistic | p.value sig |
|---|---|---|---|---|
| (Intercept) | -0.712 | 5.479 | -0.130 | 0.897 |
| MD_1000 | -0.451 | 0.085 | -5.337 | **<0.001*** ** |
| Age at MRI | 0.070 | 0.164 | 0.425 | 0.672 |
| Age at MRI^2 | -0.001 | 0.001 | -0.665 | 0.508 |
| Sex | -0.321 | 0.157 | -2.050 | **0.043*** |
| | | **adjR2**: 0.402 | **AIC**: 239.289 | **BIC**: 254.92 |

**MD_2000**

| term | estimate | std.error | statistic | p.value sig |
|---|---|---|---|---|
| (Intercept) | 0.577 | 5.409 | 0.107 | 0.915 |
| MD_2000 | -0.487 | 0.086 | -5.654 | **<0.001*** ** |
| Age at MRI | 0.025 | 0.162 | 0.155 | 0.877 |
| Age at MRI^2 | 0.000 | 0.001 | -0.363 | 0.717 |
| Sex | -0.326 | 0.154 | -2.110 | **0.037*** |
| | | **adjR2**: 0.418 | **AIC**: 236.51 | **BIC**: 252.141 |

**MA_1000**

| term | estimate | std.error | statistic | p.value sig |
|---|---|---|---|---|
| (Intercept) | -0.591 | 3.619 | -0.163 | 0.871 |
| MA_1000 | -0.779 | 0.057 | -13.711 | **<0.001*** ** |
| Age at MRI | 0.045 | 0.108 | 0.419 | 0.676 |
| Age at MRI^2 | -0.001 | 0.001 | -0.655 | 0.514 |
| Sex | 0.075 | 0.106 | 0.707 | 0.481 |
| | | **adjR2:** 0.739 | **AIC:** 156.36 | **BIC:** 171.991 |

**MA_2000**

| term | estimate | std.error | statistic | p.value sig |
|---|---|---|---|---|
| (Intercept) | 0.301 | 3.536 | 0.085 | 0.932 |
| MA_2000 | -0.787 | 0.055 | -14.196 | **<0.001*** ** |
| Age at MRI | 0.017 | 0.106 | 0.156 | 0.876 |
| Age at MRI^2 | 0.000 | 0.001 | -0.372 | 0.711 |
| Sex | -0.001 | 0.102 | -0.013 | 0.990 |
| | | **adjR2:** 0.751 | **AIC**: 151.684 | **BIC:** 167.315 |

**ODI**

| term | estimate | std.error | statistic | p.value sig |
|---|---|---|---|---|

**NDI**

| term | estimate | std.error | statistic | p.value sig |
|---|---|---|---|---|

| | | | | |
|---|---|---|---|---|
| (Intercept) | -1.670 | 4.495 | -0.371 | 0.711 |
| ODI | 0.641 | 0.068 | 9.409 | **<0.001*** ** |
| Age at MRI | 0.091 | 0.134 | 0.681 | 0.497 |
| Age at MRI^2 | -0.001 | 0.001 | -0.951 | 0.344 |
| Sex | -0.065 | 0.130 | -0.502 | 0.617 |
| | | **adjR2:** 0.597 | **AIC:** 199.67 | **BIC:** 215.301 |

| | | | | |
|---|---|---|---|---|
| (Intercept) | -0.682 | 6.227 | -0.109 | 0.913 |
| NDI | 0.083 | 0.095 | 0.873 | 0.385 |
| Age at MRI | 0.089 | 0.186 | 0.477 | 0.635 |
| Age at MRI^2 | -0.001 | 0.001 | -0.787 | 0.433 |
| Sex | -0.267 | 0.177 | -1.505 | 0.136 |
| | | **adjR2:** 0.229 | **AIC:** 264.715 | **BIC:** 280.346 |

**DCS_std**

| **term** | **estimate** | **std.error** | **statistic** | **p.value sig** |
|---|---|---|---|---|
| (Intercept) | -2.883 | 5.641 | -0.511 | 0.610 |
| DCS_std | -0.393 | 0.084 | -4.705 | **<0.001*** ** |
| Age at MRI | 0.143 | 0.168 | 0.851 | 0.397 |
| Age at MRI^2 | -0.001 | 0.001 | -1.143 | 0.256 |
| Sex | -0.158 | 0.162 | -0.978 | 0.331 |
| | | **adjR2:** 0.369 | **AIC:** 244.566 | **BIC:** 260.197 |

**DCS_skew**

| **term** | **estimate** | **std.error** | **statistic** | **p.value sig** |
|---|---|---|---|---|
| (Intercept) | -0.295 | 6.229 | -0.047 | 0.962 |
| DCS_skew | 0.104 | 0.092 | 1.129 | 0.262 |
| Age at MRI | 0.077 | 0.186 | 0.415 | 0.679 |
| Age at MRI^2 | -0.001 | 0.001 | -0.726 | 0.470 |
| Sex | -0.248 | 0.177 | -1.396 | 0.166 |
| | | **adjR2:** 0.233 | **AIC:** 264.181 | **BIC:** 279.812 |

**DCS_p75**

| **term** | **estimate** | **std.error** | **statistic** | **p.value sig** |
|---|---|---|---|---|
| (Intercept) | 0.002 | 6.090 | 0.000 | 1.000 |
| DCS_p75 | -0.211 | 0.091 | -2.318 | **0.023*** |
| Age at MRI | 0.069 | 0.182 | 0.381 | 0.704 |
| Age at MRI^2 | -0.001 | 0.001 | -0.696 | 0.488 |
| Sex | -0.375 | 0.180 | -2.087 | **0.040*** |
| | | **adjR2:** 0.264 | **AIC:** 260.012 | **BIC:** 275.643 |

**DCS_p90**

| **term** | **estimate** | **std.error** | **statistic** | **p.value sig** |
|---|---|---|---|---|
| (Intercept) | -0.058 | 6.012 | -0.010 | 0.992 |
| DCS_p90 | -0.252 | 0.090 | -2.798 | **0.006*** |
| Age at MRI | 0.070 | 0.180 | 0.387 | 0.699 |
| Age at MRI^2 | -0.001 | 0.001 | -0.698 | 0.487 |
| Sex | -0.391 | 0.177 | -2.207 | **0.030*** |
| | | **adjR2:** 0.282 | **AIC:** 257.597 | **BIC:** 273.228 |

**T1/FLAIR_ratio**

| **term** | **estimate** | **std.error** | **statistic** | **p.value sig** |
|---|---|---|---|---|
| (Intercept) | 1.858 | 6.264 | 0.297 | 0.767 |
| T1/FLAIR_ratio | 0.194 | 0.095 | 2.041 | **0.044*** |
| Age at MRI | 0.009 | 0.188 | 0.047 | 0.962 |
| Age at MRI^2 | 0.000 | 0.001 | -0.342 | 0.733 |
| Sex | -0.173 | 0.180 | -0.961 | 0.339 |

**Abbreviations:** *FA*: fractional anisotropy (b=1000, b=2000); *MD*: mean diffusivity (b=1000, b=2000); *MA*: mode of anisotropy (b=1000, b=2000); ODI: orientation dispersion index; *NDI*: neurite density index; DCS: diamagnetic component of susceptibilty (standard deviation,skewness,10th percentile, 90th percentile). All results are based on scaled variables and reported as standardized estimates. Significant results are in bold. *p<0.05; **p<0.005; ***p<0.001

**Supplementary table 2.** **Multivariate analysis for modified mode of anisotropy using DTI-ALPS$_{1000}$ as dependent variable**

**mMA_1000**

| term | estimate | std.error | statistic | p.value sig |
|---|---|---|---|---|
| (Intercept) | 0.750 | 3.480 | 0.215 | 0.830 |
| mMA_1000 | -3.265 | 0.225 | -14..539 | **<0.001*****|
| Age at MRI | 0.061 | 0.104 | 0.589 | 0.557 |
| Age at MRI^2 | -0.001 | 0.001 | -0.811 | 0.420 |
| Sex | 0.131 | 0.103 | 1.277 | 0.205 |
| | | **adjR2**: 0.759 | **AIC**: 148.42 | **BIC**: 164.05 |

**mMA_2000**

| term | estimate | std.error | statistic | p.value sig |
|---|---|---|---|---|
| (Intercept) | 3.461 | 3.303 | 1.048 | 0.297 |
| mMA_2000 | -4.220 | 0.268 | -15.721 | **<0.001*****|
| Age at MRI | 0.001 | 0.098 | 0.003 | 0.998 |
| Age at MRI^2 | -0.001 | 0.001 | -0.157 | 0.875 |
| Sex | 0.019 | 0.096 | 0.201 | 0.841 |
| | | **adjR2**: 0.784 | **AIC**: 137.38 | **BIC**: 153.01 |

**Abbreviations:** *mMA*: modified mode of anisotropy. All results are based on scaled variables and reported as standardized estimates. Significant results are in bold. *p<0.05; **p<0.005; ***p<0.001

#### Supplementary figure 1

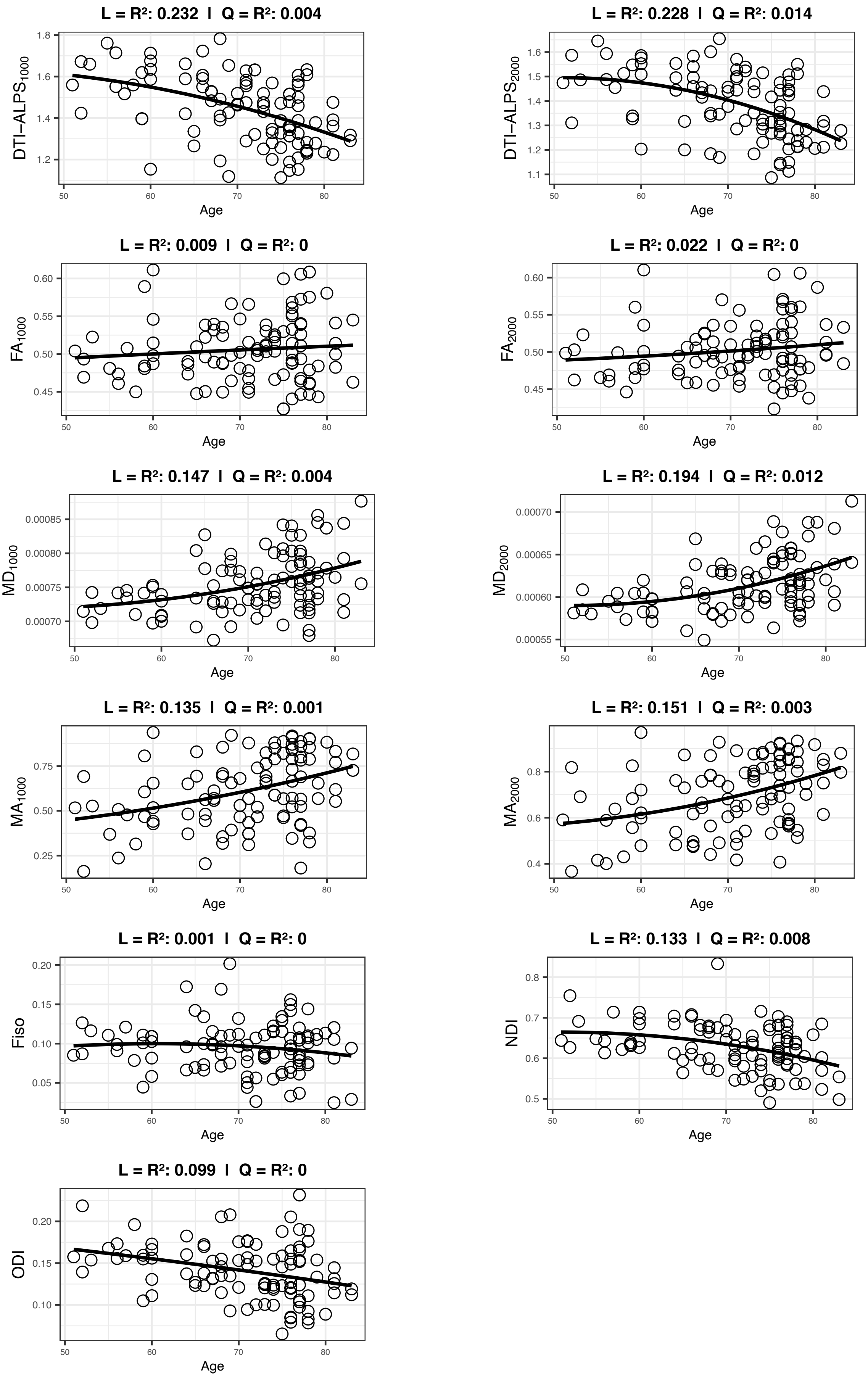

### Supplementary figure 2

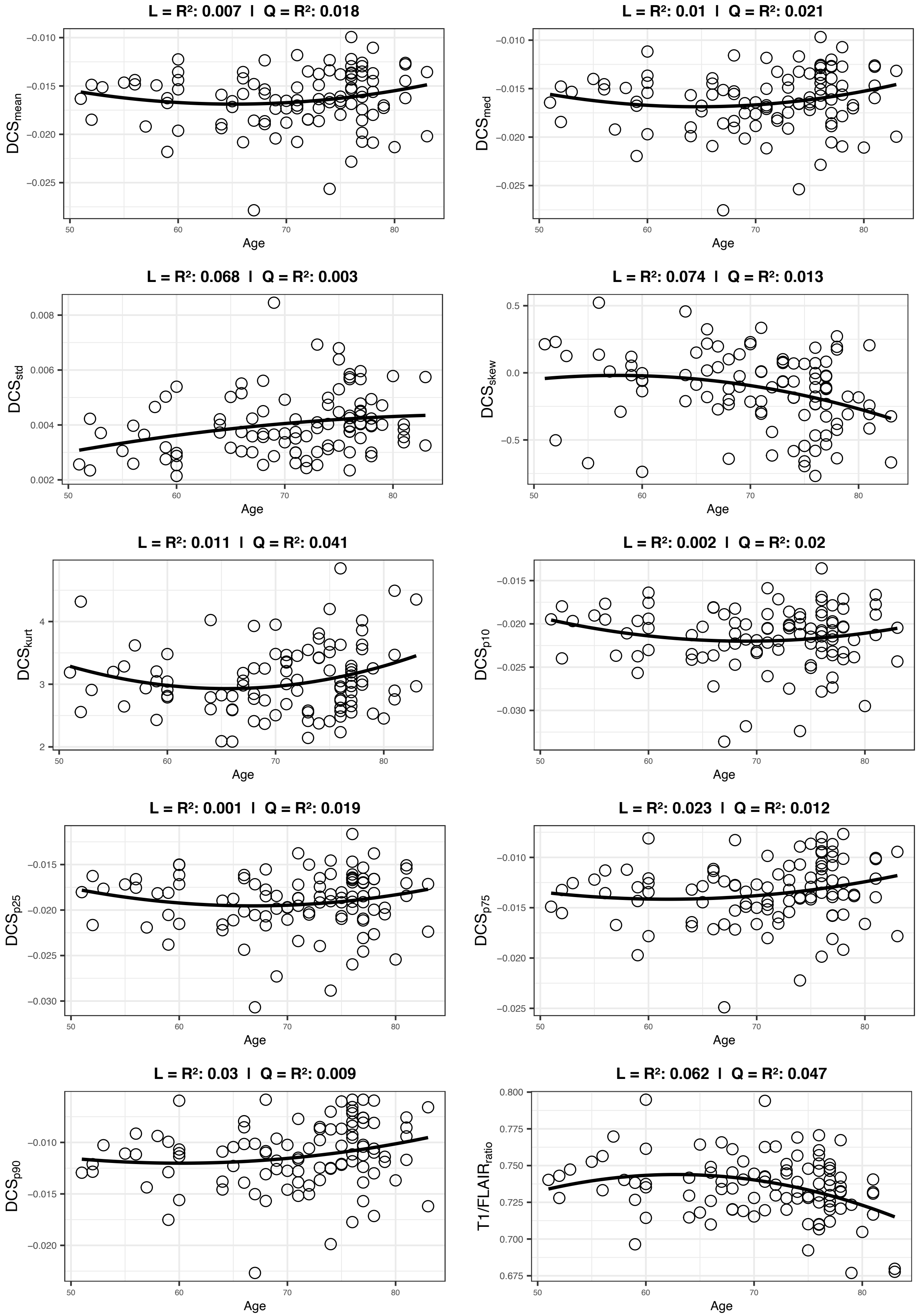
